# Targeting Neuromuscular Junction Regeneration is a Therapeutic Strategy in ALS

**DOI:** 10.1101/2025.07.13.664600

**Authors:** Samuele Negro, Chiara Baggio, Giorgia D’Este, Federico Fabris, Giulia Zanetti, Marika Tonellato, Maria Lina Massimino, Alessandro Bertoli, Roberta Schellino, Marina Boido, Monica Favagrossa, Laura Pasetto, Valentina Bonetto, Manuela Basso, Roberto Duchi, Andrea Perota, Giulia Cagnotti, Carlotta Tessarolo, Cristiano Corona, Gianni Soraru, Ilaria Genovese, Ersilia Fornetti, Alessandro Rosa, Alberto Furlan, Andrea Mattarei, Aram Megighian, Cesare Montecucco, Marco Pirazzini, Michela Rigoni

**Affiliations:** Department of Biomedical Sciences, University of Padua, Padua, Italy; U.O.C. Clinica Neurologica, Azienda Ospedale-Università Padova, Padua, Italy; Neurobiology Lab, IRCCS San Camillo Hospital, Venice, Italy; Neuroscience Institute, Section of Padua, National Research Council, Padua, Italy; Department of Neuroscience Rita Levi-Montalcini, University of Turin, Turin, Italy; Neuroscience Institute Cavalieri Ottolenghi, University of Turin, Orbassano, Italy; Research Center for ALS, Istituto di Ricerche Farmacologiche Mario Negri IRCCS, Milan, Italy; Department of Cellular, Computational and Integrative Biology (CIBIO), University of Trento, Trento, Italy; Avantea, Laboratory of Reproductive Technologies, Cremona, Italy; Department of Veterinary Sciences, University of Turin, Grugliasco, Turin, Italy; Istituto Zooprofilattico Sperimentale del Piemonte Liguria e Valle d’Aosta (IZSPLV), Turin, Italy; Department of Neurosciences, University of Padua, Padua, Italy; International Medical University UniCamillus, Rome, Italy; Center for Life Nano-& Neuro-Science, Fondazione Istituto Italiano di Tecnologia, Rome, Italy; Department of Biology and Biotechnologies “Charles Darwin”, Sapienza University of Rome, Rome, Italy; Department of Pharmaceutical and Pharmacological Sciences, University of Padua, Padua, Italy; Padua Neuroscience Center, University of Padua, Padua, Italy; Myology Center (CIR-Myo), University of Padua, Padua, Italy

**Keywords:** Amyotrophic lateral sclerosis, CXCR4, α-Latrotoxin, Neuromuscular junction, Peripheral nerve regeneration

## Abstract

Instability and denervation of the neuromuscular junction (NMJ) are early events in Amyotrophic Lateral Sclerosis (ALS), likely reflecting a progressive decline in the regenerative capacity of motor neurons (MNs) and their environment. To investigate this, we evaluated NMJ regeneration throughout disease progression in SOD1^G93A^ mice following reversible axon terminal degeneration induced by α-Latrotoxin. In parallel, we monitored the expression of CXCR4, a GPCR upregulated during axonal regeneration, and tested whether its pharmacological activation could mitigate ALS-related functional decline.

We found that NMJ regenerative capacity is largely preserved during pre- and early symptomatic stages, and remains active in subsets of NMJs even at later stages. CXCR4 is expressed at axon terminals from early disease stages, declining only at end stage. Its expression is conserved across ALS models, including SOD1^G93A^ pigs, hiPSC-derived MN with ALS mutations, and biopsies from sporadic ALS patients. CXCR4 stimulation improved motor function, NMJ innervation, MN survival, and respiratory performance in ALS mice, and axon outgrowth in iPSC-derived MN. These findings identify the NMJ and CXCR4 as viable therapeutic targets in ALS.

## Introduction

Amyotrophic lateral sclerosis (ALS) encompasses a spectrum of adult-onset diseases with multiple aetiologies, significant phenotypic heterogeneity, and incomplete understanding of its pathological mechanisms. The degeneration of upper and lower motor neurons (MNs) leads to spasticity, skeletal muscle denervation and weakness, muscle atrophy and ultimately death. ALS is mostly of sporadic origin, while ∼10% are familial forms linked to pathogenic mutations in >30 ALS-associated genes, most frequently *SOD1, TDP-43, FUS and C9ORF72*, with diverse, sometimes unclear, biological functions (Akçimen *et al*, 2023; Chia *et al*, 2018). Pathologic mutations in animal models recapitulate key aspects of human ALS and led to the identification of several aetiopathogenic mechanisms - e.g. oxidative stress, altered RNA metabolism, protein aggregation, axonal transport defects - which strongly contributed to clarifying how gene-specific alterations can harm ALS-MNs causing neurotoxicity (Hardiman *et al*, 2017; Joyce *et al*, 2011). The heterogeneity of these mechanisms hampered the identification of specific, yet common, pharmacological targets to develop therapies counteracting familial and sporadic ALS, which remains incurable (Fisher *et al*, 2023). At the same time, the neuromuscular junction (NMJ) is the initial target of pathology in both ALS patients and various animal models of the disease (Alhindi *et al*, 2022; Arbour *et al*, 2015; Chand *et al*, 2018; Clark *et al*, 2016; Fischer *et al*, 2004; So *et al*, 2018; Tremblay *et al*, 2017). This early pathological hallmark is characterized by distal muscle denervation and ensuing axonal “dying-back” degeneration, which occur before the onset of symptoms and precede MN death in the spinal cord. MN loss is asynchronous and is preceded by plastic remodelling at the NMJ, marked by cycles of denervation and reinnervation (Martineau *et al*, 2018), which continue until the definitive loss of nerve-muscle contacts and subsequent MN death. Notably, these denervation-reinnervation cycles suggest that MNs in ALS retain some regenerative competence for a period, which diminishes as the disease progresses. Consistent with this, SOD1^G93A^ MNs can regenerate their axons and reform functional synaptic connections following an acute nerve injury performed at early stages of the disease (Sharp *et al*, 2018). These findings underscore a temporal therapeutic window that could be exploited to stabilize the NMJ. Supporting or prolonging synaptic plasticity and regenerative capabilities of SOD1^G93A^ MNs at early stages, or enhancing these processes later in disease progression, could offer therapeutic benefits in ALS. However, how long ALS-MNs retain regeneration competence throughout disease progression, and whether stimulation of pro-regenerative pathways at the NMJ represent a general and realistic therapeutic strategy in ALS are presently unknown.

To assess the regeneration competence of SOD1^G93A^ NMJs throughout disease progression, we exploited α-Latrotoxin (α-LTx), which is a well-characterised presynaptic neurotoxin that induces a rapid, stereotypical, localized, synchronous and reversible degeneration of motor axon terminals (Duchen *et al*, 1981; Rigoni & Montecucco, 2017; Ushkaryov *et al*, 2008). Regenerative potential was evaluated based on neuromuscular performance and NMJ structural integrity. Additionally, we investigated the expression of CXCR4 - a marker of regenerative capability in peripheral nerves (Negro *et al*, 2017a; Negro *et al*, 2019; Zanetti *et al*) - during disease progression across different animal models of the disease, including SOD1^G93A^ pigs, in biopsies from patients, and in hiPSC-derived MNs carrying an ALS-causing pathological mutation. Finally, we explored the therapeutic potential of CXCR4 activation in SOD1^G93A^ mice to improve motor performance, NMJ innervation, respiratory function and MN survival.

## Results

### Populations of SOD1^G93A^ NMJs maintain regenerative competence throughout disease

To evaluate the regenerative competence of the neuromuscular synapse during ALS progression we exploited, as an experimental tool, the presynaptic neurotoxin α-LTx, which specifically degenerates the motor axon terminals (MAT) in a synchronous, but fully reversible manner, with complete structural and functional rescue within one week in mice (Duchen *et al*., 1981; Duregotti *et al*, 2015; Negro *et al*., 2019; Rigoni & Montecucco, 2017; Ushkaryov *et al*., 2008). Functional and anatomical regeneration were assessed by recording the evoked junctional potentials (EJPs) in individual soleus (SOL) muscle fibers of B6SJL-Tg(SOD1^G93A^) mice (hereafter referred to as SOD1^G93A^ mice) (Fig. 1), and by measuring the innervation status of the NMJs, respectively (Fig. 2). We chose the SOD1^G93A^ mouse line as to date this is the genetic model that best reproduces the decay of motor function associated with the disease in humans (Fisher *et al*., 2023). The experimental set up consists of the local injection of α-LTx in the hind limb of mutant mice of different ages that correspond to different disease stages (P30-asymptomatic, P60-early symptomatic, P90-symptomatic, P120-terminal stage) (Bilsland *et al*, 2010; Chiarotto *et al*, 2019; Tosolini *et al*, 2022; Vinsant *et al*, 2013; Zhao *et al*, 2019), and in control animals carrying the same genetic background of the mutants (B6SJL-Ntg, hereafter referred to as Ntg). EJPs were recorded 96 and 168 hours after intoxication (Fig. 1A), corresponding, respectively, to time points yielding 50% and complete functional recovery in healthy mice (D’Este *et al*, 2022; Negro *et al*, 2017b).

**Figure 1.**
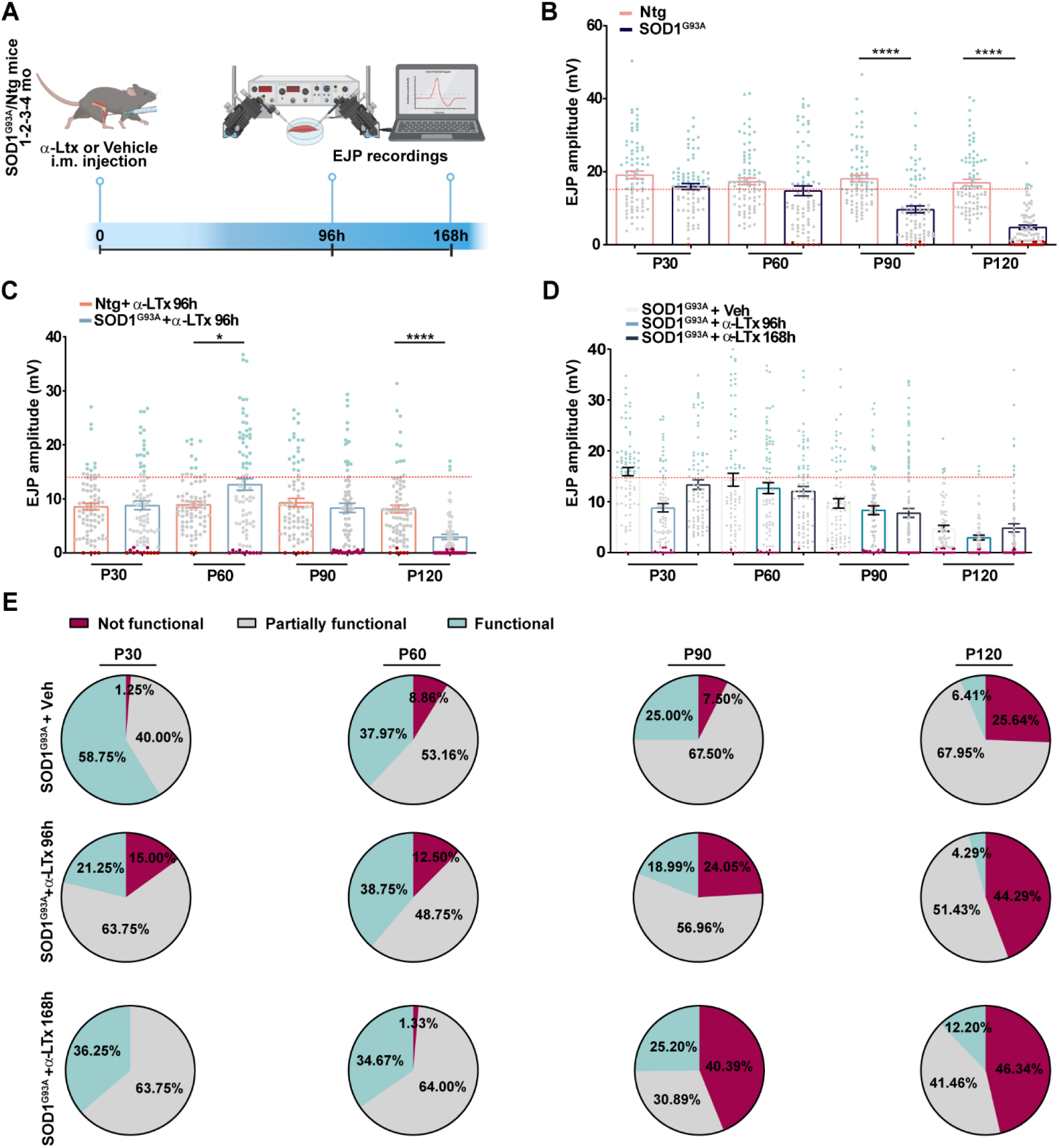
Assessment of NMJ regeneration in SOD1^G93A^ mice during disease progression. A Experimental workflow (created with Biorender). α-LTx or vehicle was i.m. injected in the hind limbs of SOD1^G93A^ and Ntg mice at different disease stages (P30, P60, P90, P120), and evoked junctional potentials (EJPs) recorded in SOL muscles 96 or 168 h later. B EJP amplitudes of SOD1^G93A^ and Ntg SOL muscle fibers over disease course. Box plots show median values and dots single-fiber distributions. Cyan dots represent fibers with EJPs above the threshold for action potential (15 mV, dotted red line), gray dots partially innervated fibers, while magenta dots fully denervated ones. N=4, 20 fibers/mouse. Ordinary One-way Anova, ****P<0,0001. C EJP amplitudes of SOD1^G93A^ and Ntg SOL fibers 96 h from α-LTx injection performed at different disease stages (P30 to P120). Box plots show median values and dots single-fiber distributions. Dots are represented using the same color-coding as in B. N=4 20 fibers/mouse. Ordinary One way Anova, *P=0,0118, ****<0,0001. D EJP amplitudes of SOD1^G93A^ SOL fibers 96 and 168h from α-LTx injection performed at different disease stages (P30 to P120). Dots are represented using the same color-coding as in B. E Pie charts depicting the proportion of fully functional (EJPs >15 mV), partially functional (1-15 mV), and fully denervated (<1 mV) NMJs in SOD1^G93A^ SOL fibers 96 and 168 h after α-LTx injection performed at different disease stages.

**Figure 2.**
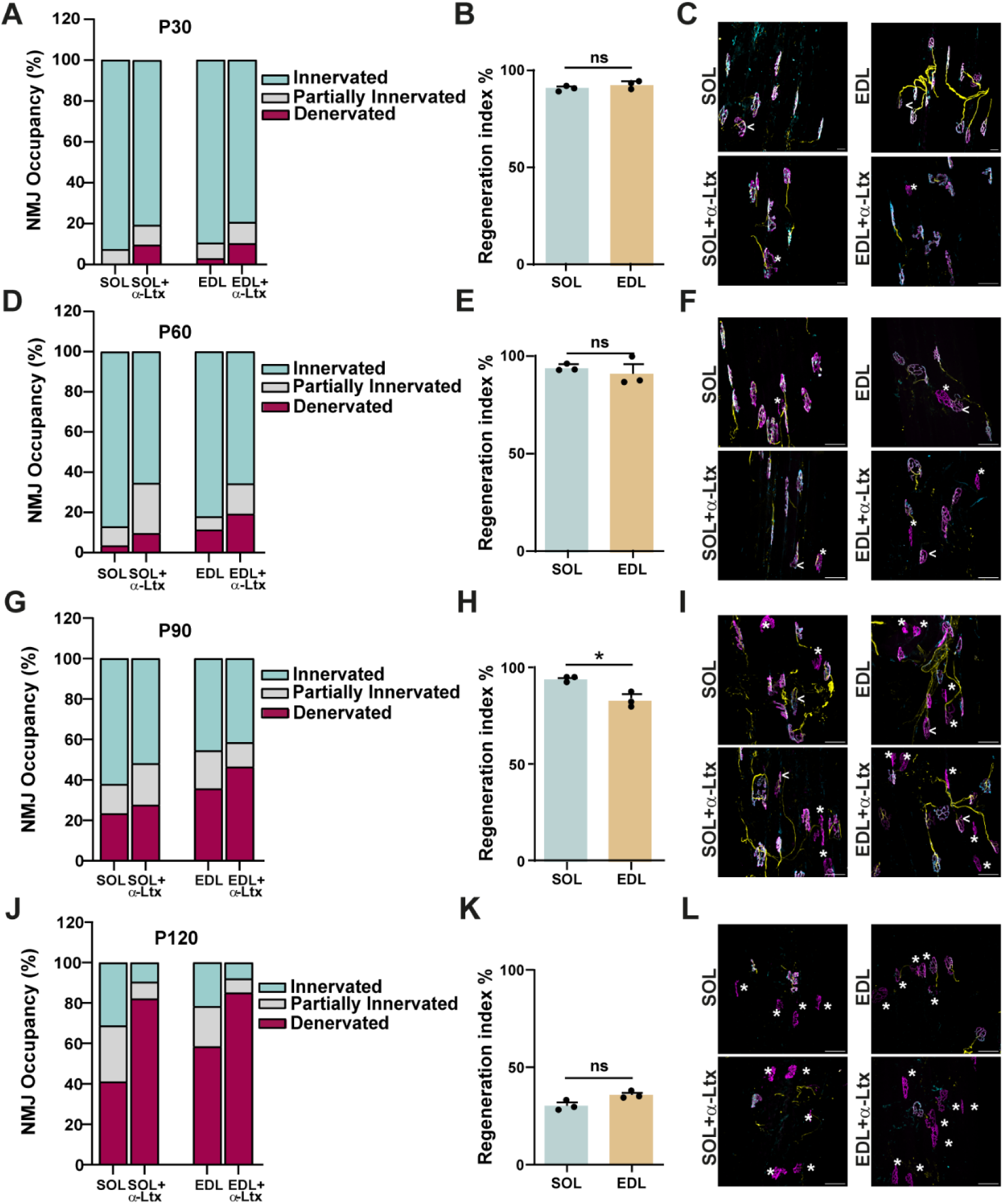
Assessment of NMJ regeneration in fast- and slow-twitch SOD1^G93A^ muscles during disease progression. A, D, G, J NMJ occupancy in SOL (slow-twitch) and EDL (fast-twitch) muscles at different disease stages (P30, P60, P90 and P120) before and 96 h after α-LTx injection. NMJs are categorized as fully innervated, partially innervated, or fully denervated based on the percentage of overlap between the presynaptic and postsynaptic compartments. B, E, H, K Regeneration index (ratio of the total number of fully and partially innervated NMJs after intoxication to those before intoxication) of SOD1^G93A^ SOL and EDL at each disease stage. Unpaired t test. *P=0,0111, ns= not significant. N=3, at least 80 NMJs analysed. Plots show median values and dots single mouse distribution. C, F, I, L Representative NMJ immunostainings in SOL and EDL 96 h post-α-LTx injection performed at the different disease stages. Presynaptic motor axon terminals are labelled for NF (yellow) and VAMP1 (cyan), while postsynaptic acetylcholine receptors are marked with fluorescent α-BTx (nAChR, magenta). White arrowheads indicate partially innervated NMJs while asterisks highlight denervated NMJs. Scale bar: 50 µm.

As a first step, we monitored neuromuscular functionality of single fibers in SOD1^G93A^ and Ntg animals over time without α-LTx injection (Fig. 1B). As expected, neurotransmission efficiency of untreated SOD1^G93A^ NMJs progressively declines over disease course compared to age-matched Ntg animals, as revealed by an increased number of fibers with EJP amplitude values below the threshold required to elicit an action potential (*i.e.*, 15 mV) (Wood & Slater, 2001) (*dotted red line*), and of fully denervated fibers (in *magenta*, each dot represents one fiber).

Next, to systematically determine the regenerative capability of SOD1^G93A^ NMJs during disease course, we injected α-LTx or vehicle in the SOL of SOD1^G93A^ and Ntg mice at different ages (spanning from pre-symptomatic to disease end-stage). α-LTx is fully active in both Ntg and SOD1^G93A^ mice, whose NMJs of different muscles undergo complete anatomical (Fig. EV1A,B) and functional degeneration of the MAT (Fig. EV1C) by 24 h. Hence, all EJPs above 1 mV recorded from 24 h onward reflect ongoing regeneration. Surprisingly, from P30 to P90, the median EJP amplitudes of SOD1^G93A^ SOL muscle fibers injected with α-LTx for 96 h at the different disease stages (*light blue boxes* in Fig. 1C) are comparable to those of age-matched intoxicated Ntg muscles (*orange boxes*) - being around half of normal values (panel B), thus corresponding to 50% functional recovery ((Negro *et al*., 2017b; Stazi *et al*, 2021), and Fig. EV1D). Only at disease end stage (P120) there is an evident reduction in neurotransmission in intoxicated mutant mice (Fig. 1C). Thus, the SOD1^G93A^ NMJs from a pre-symptomatic (P30) until a symptomatic stage (P90) recover from an acute degenerative insult in a timeframe and to an extent comparable to Ntg animals (Fig. 1C and Fig. EV1D). Strikingly, the mean EJP value of mutant NMJs at P60 is even higher than that of age-matched Ntg (Fig. 1C), indicating that the pre-symptomatic stage is endowed with the highest plasticity. The EJP amplitude distribution of single fibers for each condition reported Fig. 1C reinforces the finding that, despite the ongoing neurodegeneration (panel B), populations of mutant NMJs at P30 and P60 retain the ability to restore neurotransmission (EJP values above 15 mV, dotted line) following an acute nerve injury, with a subset of fibers at P90 showing similar recovery. These findings highlight a heterogeneous scenario in which regenerative competence of SOD1^G93A^ NMJs is largely preserved at early stages (P30-P60) and partially maintained when symptomatic (P90).

We next asked whether those fibers with an amplitude value lower than 15 mV 96 h post-toxin injection would eventually recover with additional time or instead completely fail to regenerate. We therefore recorded the EJPs 168 h after intoxication, a time point when a full rescue is achieved in healthy mice (D’Este *et al*., 2022). We found that when an acute damage is performed at P30, an extended recovery enables improved functional rescue (Fig. 1D). However, when the same injury is performed at P60 or P90, no further improvement in the mean evoked amplitude occurs upon an extended recovery period (the pre-intoxication value had almost been achieved after 96 h), rather there is a reduced number of fibres with EJP values close to zero (in *magenta* in panel D) at P60. At P120, there is a trend towards amelioration after a recovery period of 168 h.

In parallel, we calculated for each time point and experimental condition the percentage of fully functional (EJPs above 15 mV), partially functional (EJPs between 1 and 15 mV), and fully denervated (EJPs below 1 mV) NMJs, and reported the results in the pie charts in Fig. 1E. Results are in line with the findings illustrated in panel D, and confirm both the progressive impairment of neuromuscular functionality in mutant mice (top row) and that the P60 time point is endowed with the highest capability to reinnervate after transient degeneration. Indeed, 168 h after intoxication (bottom row), the percentage of fully functional fibers at P60 matches that observed before α-LTx intoxication, while the percentage of fully denervated fibers is even lower than the pre-intoxication level. Also at P90 the percentage of fully innervated fibers before toxin treatment is preserved at the end of the recovery period, meaning that a population of fibers retains the ability to recover (although lower in absolute number with respect to the P60 time point). The reduced number of partially innervated fibers, and the concomitant increase in the denervated ones at 168 h post-LTx suggest that at P90 there are pools of fibers that are unstable, part of which succumbs to the toxin’s insult. A similar scenario was found at disease end stage (P120). Nonetheless, intriguingly, there is a fraction of NMJs still able to regenerate despite the extensive degeneration characterizing this disease end-stage.

### Distinct regenerative dynamics of MN pools in SOD1^G93A^ mice

Different muscle groups, along with their innervating MNs, exhibit varying degrees of vulnerability to ALS. As fast-twitch muscle fibres are affected earlier than slow-twitch ones in the SOD1^G93A^ mouse model (Atkin *et al*, 2005; Gould *et al*, 2006; Hegedus *et al*, 2007; Kryściak *et al*, 2014; Pun *et al*, 2006; Spiller *et al*, 2016; Vinsant *et al*., 2013), we wondered whether their NMJs might display different regenerative responses to acute MAT injuries throughout the disease course. To test this, we compared the NMJ innervation status (namely NMJ occupancy) of SOL NMJs with those of the extensor digitorum longus (EDL) (predominantly innervated by slow and fast MNs, respectively) at different disease stages (from P30 to P120), before and 96 h after toxin treatment. This analysis determines the percentage of overlap between the presynaptic and postsynaptic compartments of the NMJ identified by immunofluorescence using specific markers (NF and VAMP-1 for the MAT and α-BTx for the postsynaptic specialization) (Negro *et al*, 2022) (Fig. 2). This approach confirms the progressive loss of innervation in both muscle types over time (panels A, D, G, J), which becomes more pronounced at the disease end stage (P120, panel J). SOL and EDL are differently affected by the progressive degeneration, with the EDL showing a higher number of denervated NMJs at all stages, in line with its greater vulnerability (panels A, D, G, J) (P30: SOL 0%, EDL 2.9%; P60: SOL 3.4%, EDL 11.3 %: P90: SOL 23.3%, EDL 35.6%; P120: SOL 41%, EDL 58.4%). Intoxication leads to increased denervation in the two muscles (Fig. 2 panels A, D, G), which becomes more evident at disease endpoint (panel J). We calculated the regeneration index - expressed as the ratio of the total number of fully and partially innervated NMJs after 96 h-intoxication to that before intoxication - for each disease stage. This index is nearly 100% for both muscle types at early stages (P30 and P60, panels B and E), slightly decreasing in the EDL at P90 (panel H), and dropping largely below 50% in both muscles at P120 (panel K). Of note, despite the ongoing degeneration, to which the two muscle types show different susceptibility, both the SOL and the EDL retain full regenerative competence until the symptomatic phase (P90), when this capability is slightly, though significantly, reduced in the EDL compared to the SOL. Interestingly, although the regenerative capacity progressively declines and becomes markedly compromised in the late stage (P120), a subset of NMJs appears still competent for regeneration (in line with data in Fig. 1C-E). Representative immunostainings of the innervation status of EDL and SOL for each disease stage 96 h after α-LTx injection are shown in panels C, F, I and L, where asterisks indicate denervated NMJs (lack of presynaptic staining).

We next extended the same analysis to a pure fast-twitch muscle that moves the pinna of the mouse, specifically the levator auris longus (LAL), consisting of two separate bands: a rostral (thicker, rLAL) and a caudal (thinner, cLAL) (Murray *et al*, 2010a). The rLAL has been reported to be more resistant to denervation and more sproutogenic compared to the cLAL in mice modelling spinal muscular atrophy (Murray *et al*, 2008). Fig. EV2 shows that the two bands exhibit progressive denervation (panels A, D, G), similar in extent to that of hind limb muscles (Fig. 2 panels D, G, J), more evident in cLAL than in rLAL, as expected. After transient degeneration with α-LTx, regeneration is close to 100% for both bands at P60 (panel B), consistent with values observed in the SOL and EDL. It abruptly declines at P90 (panel E), more markedly in cLAL, and then stabilizes (even increasing in cLAL) at the late-stage disease (panel H) at higher levels compared to hind limb muscles (Fig. 2).

Collectively, these data indicate that, throughout disease progression, MN pools - regardless of whether they innervate fast or slow muscles - exhibit distinct regenerative dynamics in response to nerve insults. Notwithstanding this, their regenerative capacity is largely preserved over time, peaking around P60.

### Neuronal CXCR4 expression is a common regenerative marker in ALS

The overall preservation of the regenerative potential of SOD1^G93A^ NMJs during pre-symptomatic and early symptomatic stages - and partially even after symptom onset - suggests that strategies aimed at sustaining the stability and remodeling capabilities of the NMJ could be leveraged to prevent denervation and slow disease progression. In previous studies, we have reported that CXCR4 is re-expressed by motor axon stumps following acute nerve injury, and its activation/stimulation facilitates regeneration and recovery of neurotransmission (Duregotti *et al*., 2015; Negro *et al*., 2017b; Negro *et al*., 2019; Stazi *et al*, 2020; Stazi *et al*, 2022). As CXCR4 is silenced in healthy adult peripheral nerves and reactivated under stress (Lieberam *et al*, 2005; Negro *et al*., 2017a; Negro *et al*., 2019; Zanetti *et al*, 2019), we investigated whether CXCR4 is detectable at SOD1^G93A^ NMJs during disease progression. Fig. 3 provides a representative time course of CXCR4 expression at SOD1^G93A^ SOL (panel A) and EDL (panel B) NMJs, showing that CXCR4 signal emerges early during the disease (P30) and declines as the endpoint approaches. A quantitative analysis of the percentage of CXCR4-positive NMJs at the various time points indicates that, while the peak of expression is in the early symptomatic stage, it decreases in the symptomatic phase and disappears in the late disease stage (Fig. 3C). Interestingly, no major differences in CXCR4 expression are detected between the two muscles, consistent with the overall similar regeneration index found in the SOL and EDL over the disease course (Fig. 2). Panel D illustrates the correlation between CXCR4 expression and the regenerative potential of the neuromuscular system. Specifically, the disease stage with the highest regenerative capability following toxin-induced MAT degeneration (namely P60, Fig. 1C-E) displays the highest CXCR4 expression, whereas the stage with the lowest percentage of reinnervated junctions (P120) corresponds to the lowest CXCR4 signal. These findings suggest a strong correlation between CXCR4 expression and the regenerative potential of SOD1^G93A^ neuromuscular synapses (Pearson’s coefficient r= 0.9708).

**Figure 3.**
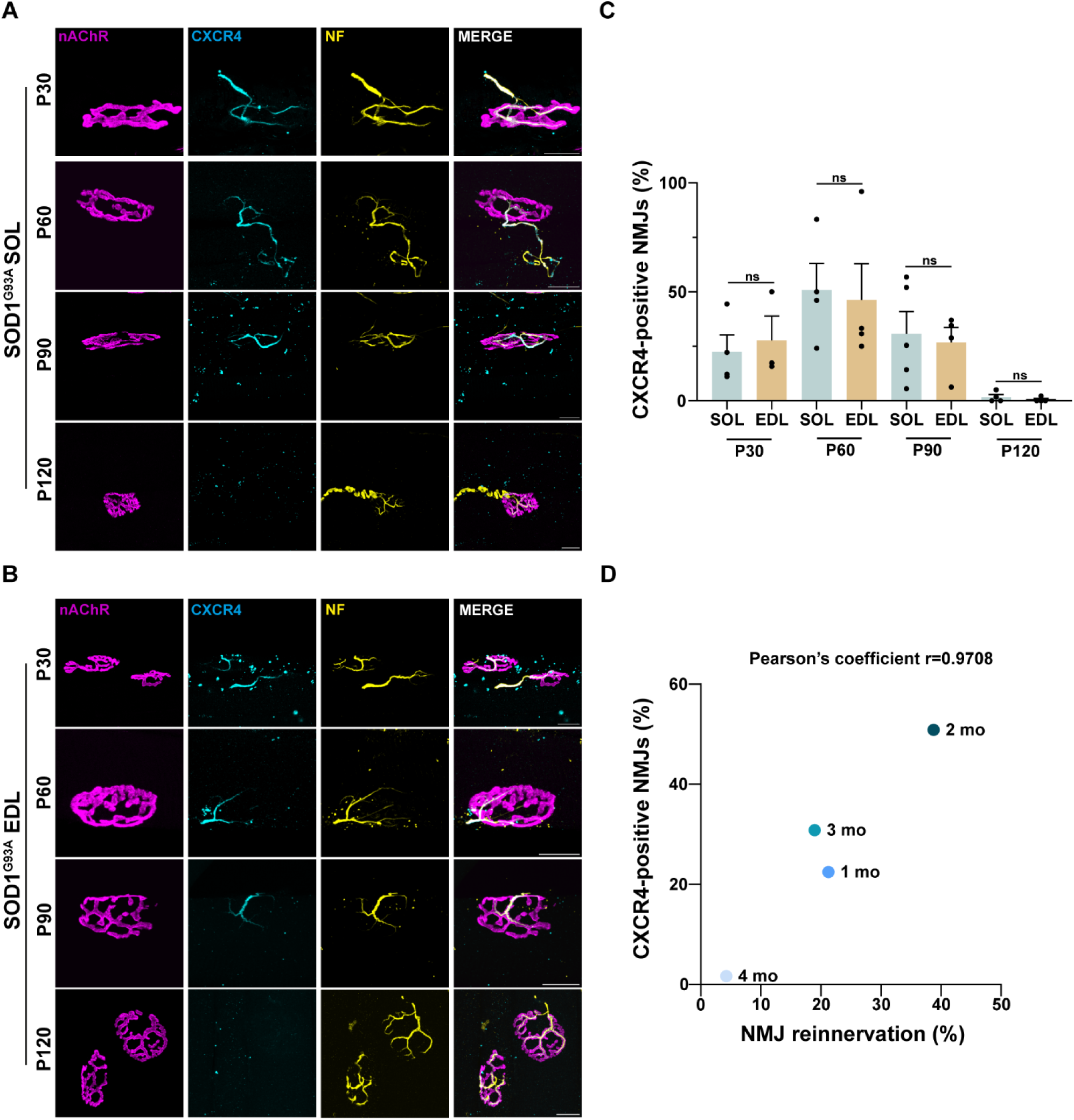
CXCR4 expression at SOD1^G93A^ NMJs correlates with NMJ regenerative potential. A-B Representative immunostainings for CXCR4 at SOL (A) and EDL (B) SOD1^G93A^ NMJs at different disease stages. Presynaptic nerve terminals are labelled for NF (yellow) and CXCR4 (cyan), postsynaptic acetylcholine receptors with fluorescent α-BTx (nAChR, magenta). Scale bar: 50 µm. C Quantification of CXCR4-positive SOD1^G93A^ NMJs at P30, P60, P90, P120. At least N=3, 100 NMJs/mouse. Plots show median values and dots single mouse distribution. D Correlation between CXCR4 expression and NMJ regenerative potential in SOD1^G93A^ mice. The highest number of regenerated NMJs following α-LTx-induced nerve terminal degeneration coincides with the peak of CXCR4 expression. Correlation test, Pearson’s coefficient r=0,9708, *P=0,0292.

Given that peripheral nerve regeneration is evolutionarily conserved in higher vertebrates and relies on mechanisms and effectors that are likely independent of specific ALS aetiologies or species, we investigated whether the MAT of several different ALS models - ranging from mice to pigs - with different disease-associated mutations, express CXCR4 in response to the progressive neurodegeneration. We analysed CXCR4 expression at the NMJs of: i) SOL and EDL from a transgenic mouse line bearing a point mutation in the TDP-43 gene (Q331K), which exhibits distal denervation, axonopathy, and MN death starting at 10 months, with minimal worsening over time (Arnold *et al*, 2013); ii) SOL and EDL from a human FUS overexpressing mouse line (hFUS^+/+^) that develops an aggressive pathology starting at 4 weeks of age, and recapitulates most aspects of ALS (Mitchell *et al*, 2013), iii) the diaphragm of a swine SOD1^G93A^ model, characterized by a prolonged phase before disease onset, with progressive denervation starting in the late pre-symptomatic stage (Crociara *et al*, 2019; Golia *et al*, 2024). CXCR4-positive NMJs were observed in all three ALS animal models starting from early disease stages (Fig. 4A-C). Additionally, we detected CXCR4 expression at the NMJ in muscle biopsies from patients carrying sALS (Table 1) (a representative staining is reported in panel D), and in human iPSC-derived MNs carrying the severe FUS pathogenic variant P525L grown in culture dishes (panel E) or microfluidic devices (MFC, panel G) (Garone *et al*, 2019). Quantification of CXCR4 signal in iPSC-derived FUS^P525L^ cultured in dishes for 12 or 20 DIV is reported in panel F. These findings point to CXCR4 as a unifying injury response marker and a sign of regenerative potential in ALS.

**Figure 4.**
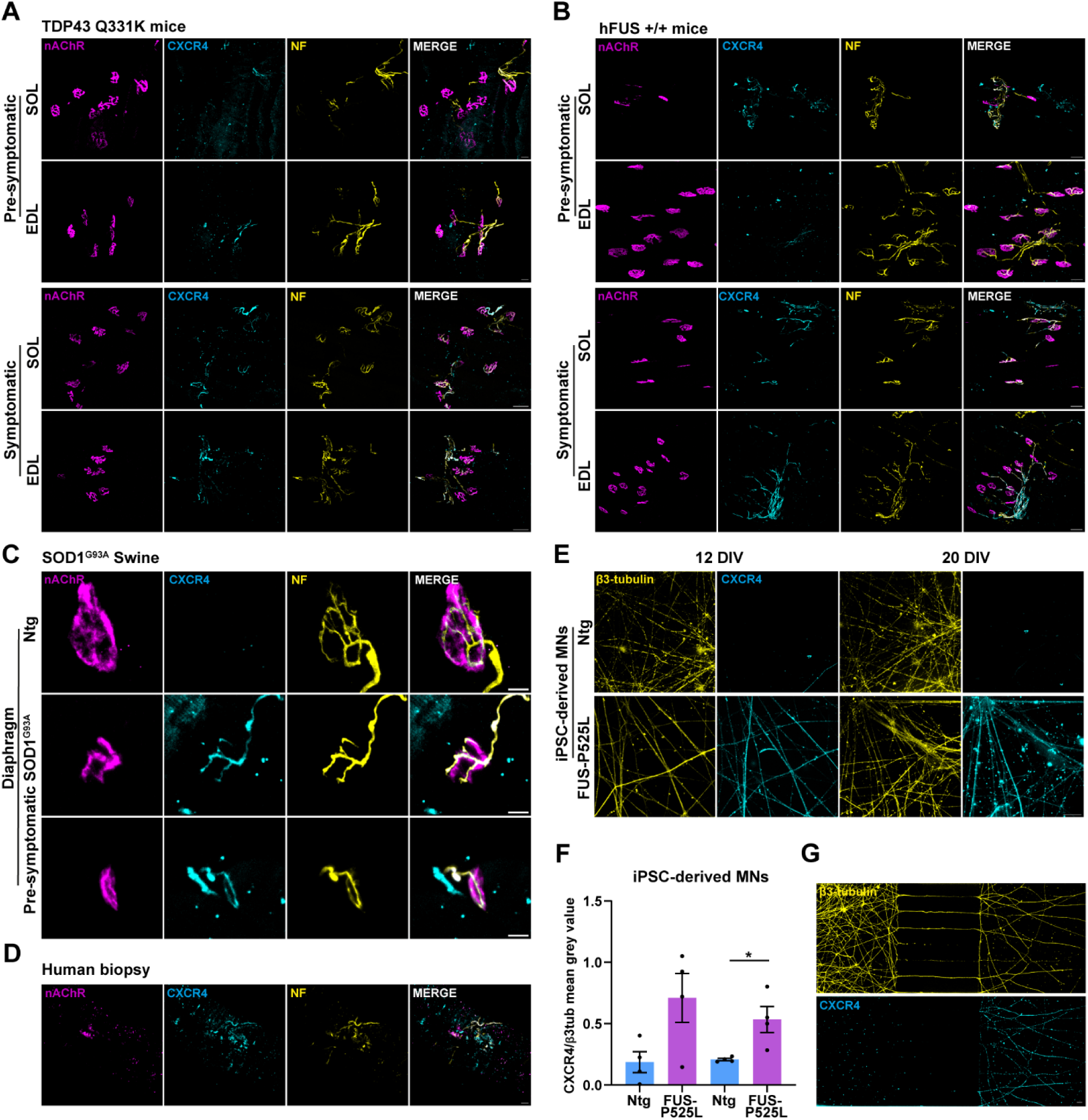
CXCR4 is a unifying injury response marker. A-C CXCR4 expression (cyan) at the NMJs of different animal models of ALS: transgenic TDP-43^Q331K^ mice (SOL and EDL) (A), hFUS-overexpressing (hFUS^+/+^) mice (SOL and EDL) (B), and SOD1^G93A^ swine model (diaphragm) (C). NMJs are labelled with fluorescent α-BTx (magenta), anti-NF (yellow) and anti-CXCR4 (cyan) antibodies. Scale bars: 50 µm. D Representative CXCR4 expression in a muscle biopsy from a sporadic ALS patient. CXCR4 is in cyan, the NMJs are identified by fluorescent α-BTx (nAChR, magenta), NF are in yellow. Scale bars: 20 µm. E, G CXCR4 expression (cyan) in human iPSC-derived MNs (β3 tubulin positive, yellow) carrying the pathogenic FUS^P525L^ mutation grown on coverslips or MFC, respectively. Scale bars: 20 µm. F Quantification of CXCR4 expression in human iPSC-derived MNs carrying the pathogenic FUS^P525L^ mutation cultured on coverslips for 12 and 20 DIV. N=3, Mann Whitney test, *P=0,0286. Plots show median values and dots single cell culture distribution.

**Table 1.** Clinical and biopsy information from human ALS patients. Demographic and clinical details of patients from whom muscle biopsies were obtained. All biopsies were taken from the vastus lateralis muscle. The table includes subject identifiers, diagnostic classification, biopsy and diagnosis dates, and the anatomical region of onset.

| ID | Gender | Age onset | Diagnosis | Date of Diagnosis | Type of onset | Site of the Onset | Site of Biopsies | Date of biopsy |
| --- | --- | --- | --- | --- | --- | --- | --- | --- |
| 24.058 | M | 54 | Sporadic | 1/10/2024 | Spinal | Upper limbs | Vastus lateralis | 31.7.24 |
| 24.023 | M | 73 | Sporadic | 12/4/24 | Spinal | Right upper limb | Vastus lateralis | 6.3.24 |

### Targeting CXCR4 delays the progressive deterioration of motor function in SOD1^G93A^ mice and protects lower MNs from cell death

Given that: i) targeting CXCR4 promotes functional recovery following acute nerve damage (Negro *et al*., 2017b; Negro *et al*., 2019; Stazi *et al*., 2020; Stazi *et al*., 2022; Stazi *et al*, 2024; Zanetti *et al*., 2019), ii) CXCR4 is expressed early by murine SOD1^G93A^ NMJs, and its expression persists throughout the disease course (Fig. 3A,B), declining only near the humane endpoint (P120), and iii) CXCR4 is consistently expressed at early stages across different ALS models, animal species and human patients, we reasoned that CXCR4 might represent a valuable target to test the outcomes of stimulating regeneration in ALS. Accordingly, we treated a cohort of SOD1^G93A^ mice with the CXCR4 agonist NUCC-390 (Mishra *et al*, 2016; Negro *et al*., 2017b; Negro *et al*., 2019; Zanetti *et al*., 2019). Treatment started at P60, a disease stage when NMJs exhibit the highest plasticity (P60) (Fig. 1C-E), which also corresponds to the peak phase of CXCR4 expression (Fig. 3C,D), and continued until P110. We assessed the impact of CXCR4 stimulation on motor performance, NMJ innervation, MN survival and the progression of axonal degeneration using a battery of readouts (Fig. 5A).

**Figure 5.**
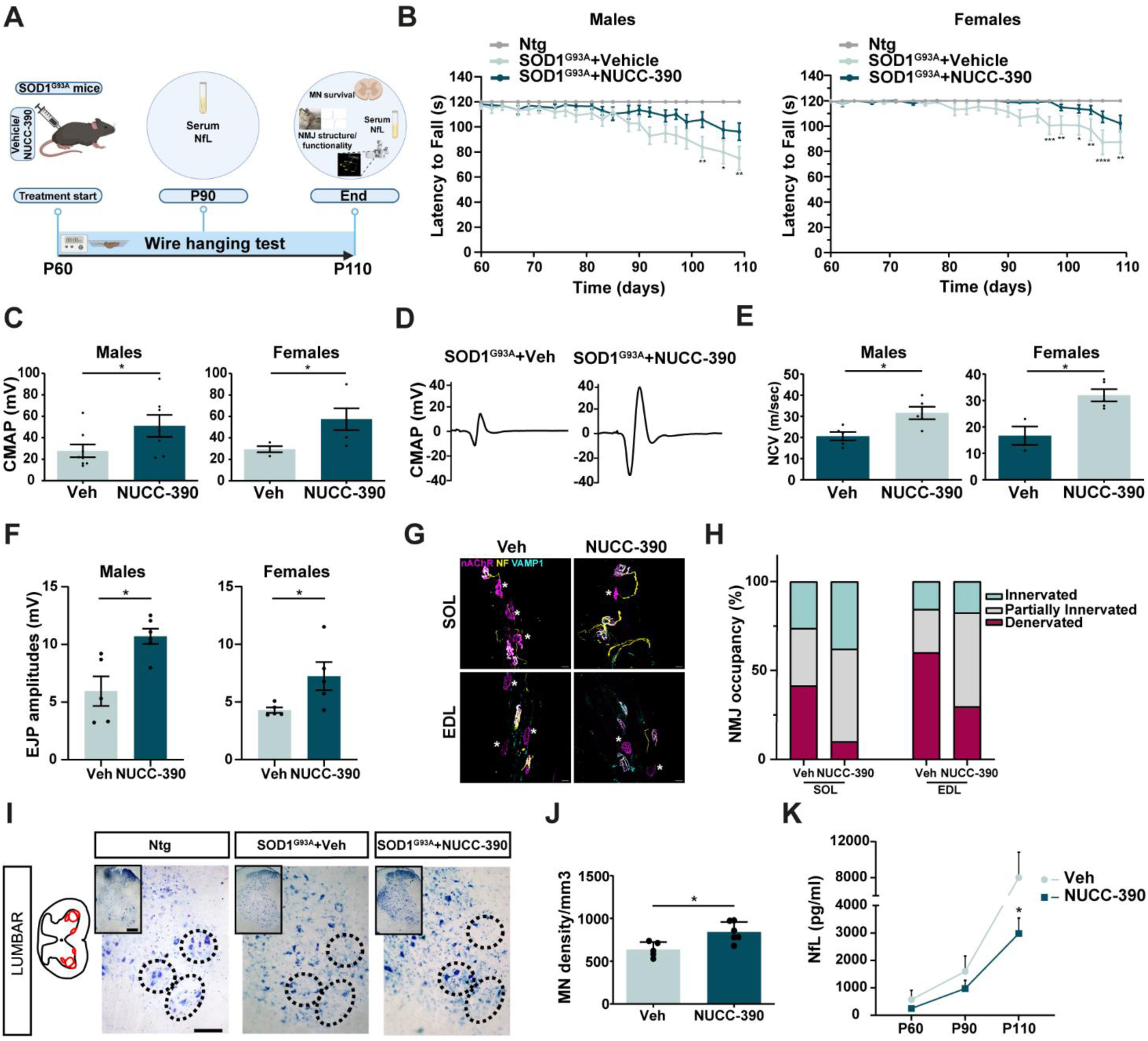
Targeting CXCR4 in SOD1^G93A^ mice delays motor function decline, improves lower MN survival and counteracts neurodegeneration. A Experimental design. SOD1^G93A^ mice were treated with the CXCR4 agonist NUCC-390 from P60 to P110. Motor function, NMJ innervation, MN survival and NfL serum levels were assessed. B Wire hanging test longitudinal assessed in SOD1^G93A^ males and females. 2 Way ANOVA, N=20 for vehicle and N=20 for NUCC-390, *P<0,05, **P<0,01, ***P<0,001 and ****P<0,0001. C CMAP recordings were performed at P110 in SOD1^G93A^ GN w/wo NUCC-390 administration started at P60. Data are expressed as CMAP amplitudes (millivolts, mV). Mann Whitney test. Females: N=4 for vehicle and N=5 for NUCC-390, *P=0,0317. Males: N=8 for vehicle and N=7 for NUCC-390, *P=0,0382. D CMAP representative traces of SOD1^G93A^ GN at P110 w/wo NUCC-390. E Nerve conduction velocity recordings performed at P110 on SOD1^G93A^ GN w/wo NUCC-390. Data are expressed in meters per second (m/s). Mann Whitney test. Females: N=3 for Veh and N=5 for NUCC-390, *P=0,0317. Males: N=5 for Veh and N=5 for NUCC-390, *P=0,0382. F EJPs recordings were performed at P110 on SOD1^G93A^ SOL w/wo NUCC-390 treatment. Data are expressed as EJP amplitudes (millivolts, mV). Mann Whitney test. Females: N=5 for Veh and N=5 for NUCC-390, *P=0,0357. Males: N=5 for Veh and N=6 for NUCC-390, *P=0,0238. G-H Representative immunostainings and quantification of NMJ occupancy at P110 in SOD1^G93A^ SOL w/wo NUCC-390 administration from P60. I Representative MN Nissl-staining of spinal cord sections at the L4-L5 level at P110 from SOD1^G93A^ mice treated or not with NUCC-390. J Quantification of lumbar MN density expressed as density of MN per mm^3^. Whitney test. N=5 for Veh and N=6 for NUCC-390, *P=0,0173. Scale bar: 100 µm, magnification: 500 µm. K NfL levels (pg/ml) measured in the blood of Veh or NUCC-390-treated SOD1^G93A^ mice from P60 till P110. Mixed effect analysis test. N=5 *P=0,0130.

A two-month treatment with NUCC-390 significantly improves motor performance at late time points, as assessed by the wire hanging assay - which evaluates the endurance of mice - both in males and females (Fig. 5B). Since endurance depends on both muscle strength and nerve functionality, we conducted electrophysiological tests to specifically assess neurotransmission in SOD1^G93A^ males and females. These included recordings of compound muscle action potentials (CMAP, panels C-D), of nerve conduction velocity (NCV, panel E) and of EJPs (panel F).

CMAP recordings performed at P110 in cohorts of SOD1^G93A^ males and females (panel C, and a representative trace in D) show a significantly slower decline in neuromuscular function in NUCC- 390 treated animals compared to SOD1^G93A^ mice treated with vehicle only, and in nerve conduction velocity (panel E). EJP amplitudes of single muscle fibres corroborate these outcomes (panel F). These findings are consistent with a higher innervation status at P110 in NUCC-390 treated SOL and EDL with respect to untreated mice (panels G,H). NUCC-390 also exerts a growth-promoting activity on iPSC-derived FUS^P525L^ MNs cultured in MFC when applied to axon terminals, indicating that also human derived neurons carrying an ALS pathological mutation are responsive to CXCR4 stimulation (Fig. EV3).

Given that distal degeneration of motor axons is likely one of the main triggers of MN cell death, we analysed MN survival in spinal cord slices from lumbar segments 4 and 5 (L4-L5) where the cell bodies of MNs innervating hind limb muscles reside. This analysis reveals a higher MN density in the ventral horns of mice that received NUCC-390, as shown by the representative staining of the correspondent spinal cord slices (Fig. 5I), and by its quantification (panel J). Moreover, NUCC- treatment leads to a reduction in the level of Neurofilament light chains (NfL) in blood samples from P110 mutant mice compared to vehicle-treated animals (panel K).

Altogether, these results support the hypothesis that targeting the periphery to enhance NMJ stability is a promising strategy to stabilize the neuromuscular synapse, thus counteracting the progressive denervation and preserving MN survival.

### Targeting CXCR4 delays the progressive decline in respiratory function of SOD1^G93A^ mice

As the respiratory function is critically affected in later stages of the disease in both mice (McCall *et al*, 2020; Romer *et al*, 2017; Stoica *et al*, 2017; Tankersley *et al*, 2007) and humans (Feldman *et al*, 2022; Lyall *et al*, 2001; Niedermeyer *et al*, 2019), we assessed whether chronic stimulation of CXCR4 by NUCC-390 could counteract the progressive impairment of this vital function. To address this, we employed a simple and sensitive assay to monitor lung ventilation (Stazi *et al*., 2022), which can be performed longitudinally on the same mouse. SOD1^G93A^ mice were treated with NUCC-390 from P60 to P110, and lung ventilation was monitored longitudinally at P60, P90 and P110 before the mice were sacrificed (Fig. 6A). Panel B shows the inferred ventilation index (IVI) of SOD1^G93A^ mice treated with NUCC-390 or vehicle, expressed as percentage of the initial value measured at P60 (before treatment start). IVI is the product of the average area of the ventilation peaks (mV × ms) and the number of peaks within 20 s. Peak areas are an estimate of the air volume inspired, directly linked to pressure variations resulting from the activity of respiratory muscles. NUCC-390 treatment starting at an early-symptomatic phase significantly slows down the progressive deterioration of the respiratory function observed from P90 to P110 in vehicle-injected mice (panel B). Panel C reports representative peak traces of vehicle and NUCC-390-treated SOD1^G93A^ mice at P60, P90 and P110, which shows a better preservation of peak amplitudes in treated mice at late time points. Ventilation measurements were complemented by the analysis of NMJ occupancy in the diaphragm (panel D-E) and MN counting at cervical (C3-C5) and thoracic (T2-T5) segments (panels F-G and H-I, respectively), where the cell bodies of MNs innervating the respiratory muscles - specifically the intercostal muscles and the diaphragm - are located. NMJ occupancy results are in line with the findings on lung ventilation, and show that CXCR4 stimulation preserves muscle innervation (panel D). A representative immunofluorescence of SOD1^G93A^ NMJs of P110 diaphragms with or without a two-month NUCC-390 treatment is reported in Fig. 6E, where asterisks identify denervated NMJs (lack of overlap between pre- and postsynaptic markers) that are more abundant in untreated samples.

**Figure 6.**
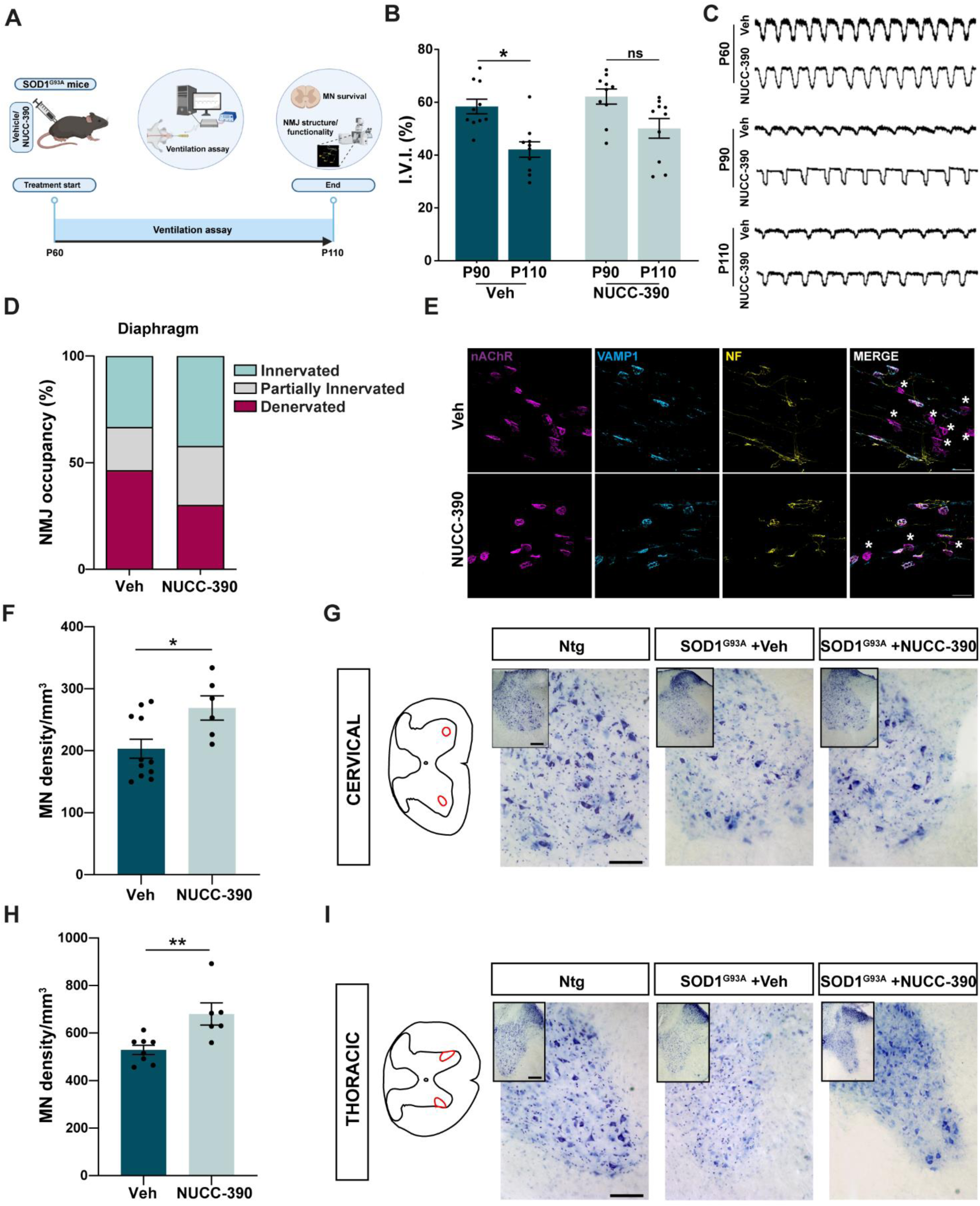
Stimulation of CXCR4 slows the progressive decline of respiratory function in SOD1^G93A^ mice. A Experimental design. SOD1^G93A^ mice were treated with NUCC-390 or Veh from P60 to P110, and lung ventilation was monitored longitudinally at P60, P90 and P110. B Inferred ventilation index (IVI) of SOD1^G93A^ mice w/wo NUCC-390. IVI is expressed as a percentage of the initial value at P60 and is calculated as the product of peak area (mV × ms), an estimate of inspired air volume, and the number of peaks in 20 sec. Kruskal-Wallis test. N=5 for both Veh and NUCC-390 groups, *P=0,0287, ns: not significant. C Representative ventilation peak traces of SOD1^G93A^ mice w/wo NUCC-390 measured at P60, P90 and P110. D Quantification of NMJ occupancy at P110 in SOD1^G93A^ diaphragms w/wo NUCC-390. E Representative NMJ immunostainings of SOD1^G93A^ diaphragms at P110. Presynaptic nerve terminals are labelled for VAMP1 (cyan) and NF (yellow), postsynaptic acetylcholine receptors are marked with α-BTx (nAChR, magenta). Asterisks indicate denervated NMJs. Scale bar: 50 µm. F-I Analysis of MN survival in cervical (C3-C5) and thoracic (T2-T5) spinal cord levels labelled by Nissl staining (blue) at P110 in SOD1^G93A^ mice w/wo NUCC-390 treatment (from P60). MN survival is expressed as density of MN/mm^3^ (F, H). Representative images (G, I). Mann-Whitney test. F: N=11 for Veh and N=6 for NUCC-390, *P=0,0273. H: N=8 for Veh and N=6 for NUCC-390, **P=0,0080.

CXCR4 engagement allows a higher survival rate of cervical and thoracic MNs (Fig. 6 F-I), a result consistent with the increased MN survival in the lumbar spinal cord (Fig. 5I,J).

Collectively, these data show that CXCR4 stimulation counteracts the progression of respiratory dysfunction in SOD1^G93A^ mice by preserving NMJ innervation and mitigating MN death in the spinal cord.

## Discussion

In this study, we systematically characterized the regenerative capacity of SOD1^G93A^ NMJs during disease progression by analysing neurotransmission recovery in individual muscle fibres and NMJ innervation following an acute presynaptic insult. We found that NMJs in SOD1^G93A^ ALS mice possess a strong and previously underappreciated regenerative capacity during early and mid-disease stages. A key strength of the study is the identification of CXCR4 as a functional biomarker and a target to improve regeneration. By pharmacologically activating CXCR4 with an agonist compound, we achieved significant improvements in MN survival, NMJ integrity and respiratory function, and a reduced axonal degeneration.

Our research builds on experimental and clinical evidence appointing the neuromuscular synapse as primary site of degeneration in ALS (Clark *et al*., 2016; Dadon-Nachum *et al*, 2011; Fischer *et al*., 2004; Frey *et al*, 2000; Picchiarelli *et al*, 2019; Rocha *et al*, 2013; Wishart *et al*, 2006), as well as on data from animal models suggesting a transient window of NMJ plasticity before degeneration becomes predominant (Martineau *et al*., 2018; Sharp *et al*., 2018). These observations suggest that promoting NMJ plasticity may help to slow progressive denervation, enhance MN survival, and preserve muscle responsiveness to reinnervation.

We have developed an innovative approach to identify the molecular drivers of nerve terminal regeneration at the murine NMJ using the presynaptic neurotoxin α-LTx (Duchen *et al*., 1981; Rigoni & Montecucco, 2017; Ushkaryov *et al*., 2008). This toxin induces a calcium-mediated, synchronous and reversible degeneration of the MAT, which fully recovers anatomically and functionally within one week in mice (D’Este *et al*., 2022; Duregotti *et al*., 2015; Duregotti *et al*, 2013; Negro *et al*., 2017b; Rigoni *et al*, 2007). Here, we exploited this toxin as an experimental tool to identify the window of regeneration competence of single SOD1^G93A^ fibres during disease progression. Indeed, nerve terminal degeneration and regeneration induced by the neurotoxin mimics the remodelling of mutant NMJs during the initial stages of the disease but occur in a synchronous manner, allowing for the quantitative analysis of reinnervation through electrophysiology coupled to imaging. We injected α-LTx in the hind limbs of SOD1^G93A^ mice at pre-symptomatic, early-symptomatic and symptomatic disease stages (Bilsland *et al*., 2010; Chiarotto *et al*., 2019; Vinsant *et al*., 2013; Zhao *et al*., 2019) and recorded the EJPs of single fibres of SOL at 96 and 168 h post-injection, corresponding to half and full recovery in intoxicated wild-type mice, respectively (D’Este *et al*., 2022). This analysis highlighted a heterogeneous scenario in which regenerative competence at SOD1^G93A^ NMJs is preserved at early stages (maximally at P60) and partially maintained at symptomatic ones - despite the ongoing neurodegeneration - a finding also supported by a parallel analysis of NMJ innervation. These results align with previous observations that an acute injury (by nerve compression) to SOD1^G93A^ motor nerves performed at early disease stages promotes muscle reinnervation and survival of functionally intact nerve-muscle contacts (Sharp *et al*., 2018). This suggests that sustaining the plasticity of the neuromuscular system could hold therapeutic potential.

Comparison of denervation/reinnervation rates at SOD1^G93A^ NMJs of different muscles shows that MN pools do not respond uniformly to chronic insult - neither in terms of timing nor in terms of severity of degeneration. Nonetheless, a shared regenerative capacity upon the toxin insult is observed - albeit varying in timing and extent across different pools - peaking around P60. This result points to this early symptomatic stage as a critical window during which NMJs are most responsive to reinnervation and pro-regenerative stimuli. In addition, our findings indicate that heterogeneity in the injury response exists also within more anatomically and functionally uniform regions (Murray *et al*., 2008; Murray *et al*, 2010b). For instance, even within a muscle like the LAL, which is composed of a relatively homogeneous population of fast-twitch muscle fibres, individual MNs display differential susceptibility to degeneration, and as well retain distinct regeneration/resilience properties.

Our recent work revealed the G-protein coupled receptor CXCR4 as a marker of reparative competence, and the CXCR4-CXCL12α axis as a crucial driver of nerve repair. Indeed the receptor, highly expressed in the peripheral nervous system during development (Bonanomi *et al*, 2019; Lieberam *et al*., 2005) but absent in the adult, is re-expressed upon acute peripheral nerve injuries by the regenerating motor axon stump, and its engagement by its natural ligand CXCL12α or an agonist molecule promotes a faster recovery of function (Negro *et al*., 2017b; Negro *et al*., 2019; Rigoni & Negro, 2020; Stazi *et al*., 2020; Stazi *et al*., 2022; Stazi *et al*., 2024). This finding suggests that signals produced at the injury site trigger CXCR4 re-expression, and that CXCR4 represents a valuable pharmacological target for enhancing the regenerative ability of peripheral neurons.

Here, we found that CXCR4 is consistently expressed at NMJs from the early stages of ALS across different animal models of the disease, regardless of the aetiology, in human biopsies and in iPSC-derived MNs carrying a ALS-related pathogenic mutation (Silvestri *et al*, 2024) thus representing a unifying potential pharmacological target across multiple forms of ALS. Noticeably, the NMJs of predominantly fast- (EDL) and slow-twitch (SOL) muscles express the receptor at similar times and levels. Hence, despite their differing vulnerabilities to neurodegeneration (Atkin *et al*., 2005; Gould *et al*., 2006; Hegedus *et al*., 2007; Kryściak *et al*., 2014; Pun *et al*., 2006; Ragagnin *et al*, 2019; Spiller *et al*., 2016; Vinsant *et al*., 2013), SOL and EDL NMJs exhibit an overall comparable regenerative capability, which declines in both muscles during the final stages of the disease, and which could be targeted by CXCR4 activators to support plasticity of the system. Importantly, CXCR4 expression occurs before denervation in both murine (SOL and EDL) and swine (diaphragm) muscles (Golia *et al*., 2024), putatively identifying this receptor as an early stress marker.

Treatment of a cohort of SOD1^G93A^ mice with a CXCR4 agonist starting at the disease stage characterized by the maximum regenerative capacity (P60) leads to a slower decline of motor performance and respiratory function in the late stages, preservation of NMJ innervation, and reduced MN loss and axonal degeneration.

These findings indicate that enhancing the intrinsic regenerative capacity of the peripheral nervous system is a promising strategy to counteract denervation, preserve muscle responsiveness to reinnervation, and support MN survival in ALS. By promoting regeneration of the NMJ, it is possible to sustain neuromuscular function, maintain the potential for reinnervation, and prolong both motor neuron viability and respiratory performance. These results lay the groundwork for future combinatorial therapies aimed at engaging endogenous repair mechanisms through pharmacological targeting of peripheral axons, with the goal of slowing disease progression in ALS.

## Methods

### Ethical statement

Animals were housed with controlled photoperiod and temperature, with free access to standard food and water. Mice were handled by specialised personnel under the control of inspectors from the Veterinary Service of the Local Sanitary Service (ASL 16-Padua), who are the local officers of the Ministry of Health. The use of animals and the experimental protocols were approved by the Ethical Committee and by the Animal Welfare Coordinator of the “Organismo Preposto al Benessere Animale” of the University of Padua. All procedures are specified in the projects approved by the Italian Ministry of Health, Ufficio VI (526/2019 PR and 922/2021 PR), and were conducted in accordance with national laws and policies (D.L. n. 26, March 14, 2014), and following the guidelines established by the European Community Council Directive (2010/63/EU) for the care and use of animals for scientific purposes.

### Experimental models

#### Murine models

The B6SJL-Tg(SOD1^G93A^)1Gur/J transgenic mice line - hereafter referred to as SOD1^G93A^ mice - expressing the mutated form of human SOD1 with a transition from a Glycine to an Alanine in position 93 (G93A) (Gurney *et al*, 1994) was purchased from Jackson Laboratories (https://www.jax.org/jax-mice-and-services/preclinical-research-services/neurobiology-services/amyotrophic-lateral-sclerosis-efficacy-studies/als-mouse-model-resource, cat. n. 002726). The strain was maintained by breeding hemizygous carriers B6SJLTg(SOD1^G93A^)1Gur/J (males) with a hybrid B6SJLF1 female. B6SJLF1 females were obtained by breeding SJL/J male mice with C57BL/GJ, hence obtaining a hybrid F1 generation. B6SJL-Tg(SOD1^G93A^)1Gur/J line carries a 40% expansion in transgene copy number and a 4-fold increase in SOD1 activity. Genotyping was performed via standard PCR (MyTaq Extract-PCR kit, Bioline) with supplied primers (Jackson Laboratory). Non-transgenic (B6SJL-Ntg) - hereafter referred to as Ntg mice - littermates were used as a negative control.

The TDP-43^Q331K^ (C57BL/6NJ background) transgenic mice line displays a point mutation in the TDP- 43 gene that causes a transition from Glutamine to Lysine, and undergoes a slow progressing mild neuropathy characterised by distal denervation, axonopathy and MN death starting from 10 months (Arnold *et al*., 2013). Muscle samples from transgenic animals of 3 and 10 months of age (TDP- 43^Q331K^ mice) and their respective controls (TDP-43^WT^) were kindly provided by Prof. M. Basso (Dept. of Cellular, Computational and Integrative Biology (CIBIO), University of Trento, Italy).

In the PrP-hFUSWT3Cshw/J mice line, the PrP-hFUS transgene contains a mouse prion protein promoter directing expression of a hemagglutinin-tagged human wild type ‘fused in sarcoma cDNA’ sequence (HA-hFUS). Homozygous PrP-hFUS mice from founder line WT3 (PrP-hFUS-WT3) have FUS overexpression levels that result in an aggressive neurodegenerative phenotype recapitulating many features of human ALS (Mitchell *et al*., 2013). Significant impairment and decline in rotarod performance start from 4 weeks of age, with reduced locomotor activity and abnormal hind limb splay from 6 weeks of age. Muscle samples from transgenic animals (hFUS^+/+^ mice) at pre-symptomatic and symptomatic phases were kindly provided by Dr. V. Bonetto (Istituto di Ricerche Farmacologiche Mario Negri IRCCS, Milan, Italy).

#### Swine model

The hSOD1^G93A^ transgenic swine model expressing the human pathological hSOD1^G93A^ allele (Crociara *et al*., 2019) was generated by Avantea (Cremona), Authorization n° 1187/2020-PR by the Italian Ministry of Health according to Dgls 26_2014. Snap-frozen samples of hSOD1^G93A^ swine diaphragms at presymptomatic stage were collected after euthanasia at Avantea and kindly provided by Dr. C. Corona (Istituto Zooprofilattico Sperimentale del Piemonte Liguria e Valle d’Aosta IZSPLV, Turin, Italy).

#### Human biopsies

The Neuromuscular Bank of Tissues and DNA samples, member of the Telethon Network of Genetic Biobanks (project no. GTB12001), funded by Telethon Italy, and of the EuroBioBank network, provided us with specimens (Table 1) through Prof. G. Sorarù, Dept. of Neurosciences, University of Padua, Italy). The project was approved by Comitato Etico per la Sperimentazione Clinica della Provincia di Padova on 17th December 2020, CESC code: 4988/A0/20 and URC code: AOP2208.

#### hiPSC-derived FUS^P525L^MNs

The wild-type (WT) iPSC line WTSIi004-A was obtained from the European Bank for induced Pluripotent Stem Cells (EBiSC; hPSCreg name: WTSIi004-A, RRID:CVCL_AI02; as stated by EBiSC, informed consent was obtained from the donor, NRES Committee Yorkshire & The Humber-Leeds West approval number 15/YH/0391). The isogenic WTSI- FUS^P525L^ iPSC line was derived from these cells using TALEN-directed mutagenesis (Silvestri *et al*., 2024). Stable and inducible iPSCs for spinal MN differentiation (NIL iPSCs) were generated following an established protocol (Garone *et al*., 2019). Mature MNs were maintained in Neurobasal/B27 medium, supplemented with 20 ng/mL L-ascorbic acid (Merck Life Sciences), 20 ng/mL BDNF (PreproTech), and 10 ng/mL GDNF (PreproTech) by replacing culture medium every other day until either day 12 or 20. Thereafter, cells were fixed and processed for immunofluorescence (Silvestri *et al*., 2024).

SOD1^G93A^ mice were employed for behavioural tests, electrophysiological recordings, immunostainings, and assessment of the respiratory function, MN survival and Neurofilament light chains levels in the blood (w/wo chronic treatments). Soleus (SOL) and extensor digitorum longus (EDL) muscles from TDP-43^Q331K^ and hFUS^+/+^ mice, diaphragms from SOD1^G93A^ swines, human muscle biopsies from patients carrying ALS, and human iPSC-derived MNs (carrying the pathogenic FUS^P525L^ mutation) were used for CXCR4 immunodetection.

### Reagents

The following primary/secondary antibodies/fluorescent conjugates were employed: rabbit polyclonal anti-vesicle associated membrane protein 1 (VAMP-1) (generated as described in (Rossetto *et al*, 1996), 1:200), chicken polyclonal anti-Neurofilament (NF) (Abcam, cat. Ab 4680, 1:1000), rabbit polyclonal anti-CXCR4 (Abcam, cat. Ab 124824, 1:200), α-Bungarotoxin Alexa555- conjugated (α-BTx-555) (Life Technologies, cat. B35451, 1:200). Secondary antibodies Alexa-conjugated (1:200) were from Life Technologies. α-Latrotoxin (α-LTx) was purchased from Alomone (cat. LSP-130). The CXCR4 agonist NUCC-390 was synthesised as in (Negro *et al*., 2019). Unless otherwise stated, all other reagents were from Sigma.

### α-LTx and NUCC-390 in vivo administration

Upon isoflurane anesthetization, mice were intramuscularly injected in the hind limb with α-LTx (5 μg/kg) diluted in 15 μl of physiological saline (0.9% w/v NaCl in distilled water) as previously described (Duregotti *et al*., 2015; Negro *et al*., 2017b). Control animals were injected with saline.

The pharmacological protocol employed for treatment with the CXCR4 agonist NUCC-390 is based on our previous work (Negro *et al*., 2019). Mice received the compound i.p. 3 times a week at a concentration of 3.2 mg/Kg (diluted in 0.9% NaCl, 0.2% gelatin solution, 15 µl injection volume). Injections were performed approximately at the same hour of the day. Weight was constantly monitored to meet the ethical guidelines.

### Motor function assessment

Assessment of motor functionality was performed in Ntg and SOD1^G93A^ mice by wire hanging test and electrophysiology.

#### Wire hanging test

Performed in SOD1^G93A^ mice 3 times a week, starting one week before the first treatment (veh/NUCC-390 started at post-natal day 60, P60) for training, until P110. The test assesses the ability of a mouse to hang when suspended on a flat grid. The animal is placed on a flat grid balanced on a box, 30 cm above the floor (coated with a soft structure to avoid injuries). The grid is turned carefully, paying attention to flip the animal backward in order for it to hold the grip. The time the mouse can hang with all the limbs is monitored, and the test is terminated if the animal releases the grid with its hind legs for more than 10 seconds or falls off. The test is repeated 3 times with at least 60 sec between every repetition to allow recovery. The latency to fall, which represents the longest time the mouse can hang on the grid (max 120 sec), is recorded; if the maximum score is achieved the test is not repeated. Training of the animals is performed for three consecutive days before the first recording, which is executed at the same hour of the day. The best score of every session is recorded for each mouse and these values are pooled to obtain the mean for each time point.

#### Electrophysiology

Evoked Junctional Potential (EJP) recordings were performed in SOL of Ntg or mutant mice. Anesthetized mice were locally injected at different ages/disease stages (P30, P60, P90, P120) with α-LTx (5 μg/kg in 0.9% NaCl, 0.2% gelatin solution) or vehicle (0.9% NaCl, 0.2% gelatin solution) using a 701 N Hamilton® syringe (Merck, 20,779), and muscles collected for analysis 96 or 168 h later. Contralateral, vehicle-injected muscles were used as internal controls. In another set of experiments, EJPs were recorded at P110 in vehicle or NUCC-390-treated SOD1^G93A^ mice (from P60).

EJPs were recorded as previously described (Zanetti *et al*, 2018). Briefly, SOL were dissected and immediately placed in an oxygenated (95% O_2_ and 5% CO_2_) experimental chamber for recording. Intracellular recordings of EJPs were performed in Krebs-Ringer solution at RT (20-22°C) using intracellular glass microelectrodes (1.5 mm outer diameter, 1.0 mm inner diameter, 15-20 MΩ tip resistance; GB150TF, Science Products GmbH Germany), filled with a 1:2 solution of 3 M KCl and 3 M CH_3_COOK. Evoked neurotransmitter release was recorded in current-clamp mode, and resting membrane potential was adjusted with current injection to −70 mV. EJPs were elicited by supramaximal nerve stimulation at 0.5 Hz using a suction glass microelectrode (GB150TF, Science Products GmbH Germany) connected to a S88 stimulator (Grass, USA). Muscle fibre contraction during intracellular recordings was blocked by adding 1 μM μ-Conotoxin GIIIB (Alomone Lab, Israel). Intracellularly recorded signals were amplified with an intracellular amplifier (BA-01X, NPI, Germany). Using a digital A/C interface (NI PCI-6221, National Instruments, USA), amplified signals were converted to digital format offline using WinEDR (Strathclyde University; pClamp, Axon, USA). Stored data were analyzed off-line using the software pClamp (Axon, USA). At the end of the experiment, muscles were processed for indirect immunofluorescence.

Compound muscle action potential (CMAP) recordings were performed at P110 in anesthetized SOD1^G93A^ gastrocnemius muscles (GN), w/wo NUCC-treatment starting from P60. The sciatic nerve was exposed under general anaesthesia (Avertin, final concentration 0,25mg/ml, i.p. injection) through an incision in the trochanteric region by removing the skin and biceps femoris muscle. Curved forceps were used to gently disrupt the connective tissue beneath the nerve, allowing the placement of a 1 cm-wide parafilm strip underneath as a reference marker. A pair of stimulating needle electrodes (Grass, USA) were positioned on the exposed nerve using a mechanical micromanipulator (MM33, FST, Germany). Electromyographic activity of the GN was recorded using two needle electrodes (Grass, USA): the recording electrode was inserted halfway into the muscle, while the indifferent electrode was placed in its distal tendon. CMAPs were recorded following supramaximal stimulation of the nerve at 0.5 Hz (0.4 ms stimulus duration) using a stimulator (S88, Grass, USA) via a stimulus isolation unit (SIU5, Grass, USA) in capacitance coupling mode. Stimulus intensity was progressively increased until CMAP amplitude plateaued (5–15 mV in controls; up to 50 mV after nerve injury). Recorded signals were amplified (P6, Grass, USA), digitized (National Instruments, USA), and analysed using WinEDR (Strathclyde University) and pClamp (Axon, USA). CMAP amplitudes were calculated and expressed in mV. At the end of the experiment, GN muscles were processed for immunofluorescence.

Nerve conduction velocity (NCV) was performed at P110 in SOD1^G93A^ mice w/wo NUCC-treatment starting at P60. NCV was determined under general anaesthesia (Avertin, final concentration 0,25mg/ml, i.p. injection) by inserting a recording electrode halfway into the GN, stimulating the sciatic nerve at two distinct sites, and measuring (a) the latency difference between CMAPs, and (b) the inter-electrode distance. Latency was defined as the interval between nerve stimulation and the onset of the CMAP wave. The 1 cm parafilm strip served as a reference for stimulation sites. NCV was expressed in m/sec.

### Lung ventilation assessment

Lung ventilation was measured by an indirect, highly sensitive test. A probe connected to a pressure sensor was placed inside the oesophagus at the mediastinum level in anaesthetised mice. Asymmetric peaks that match the frequency of lung ventilation events characterise the signal pattern we recorded. The peak area may be taken as an estimate of the volume of air inspired, which is directly related to pressure variations resulting from the activity of muscles involved in breathing.

Following general anaesthesia (a cocktail of Ketamine 6,25 mg/kg and Xylazine 2,5 mg/kg), animals were left 10 min in their cages to relax. Two-month-old SOD1^G93A^ mice were i.p. injected every 2 days with NUCC-390 (diluted in 40 μl PBS containing 0.2% gelatin) or vehicle until P110. For each mouse, recordings were performed longitudinally at the day of the first treatment (P60) and repeated at P90 and P110. A 20 ga x 38 mm plastic feeding tube (Instech Laboratories, Inc.) attached to a pressure sensor (Honeywell, 142PC01D) was carefully positioned in the mice oral cavity and placed in the lower third of the oesophagus at the level of the mediastinum. Animals were laid on their left side on a pre-warmed surface to record their lung pressure variations, which were used to infer animal ventilation by recording signals that were amplified and digitised with WinEDR V3.4.6 software (Strathclyde University, Scotland). Stored data were analysed using Clampfit software (Axon, USA). We recorded 2 min ventilation for each animal to ensure trace stability, and then 20 sec of each trace were analysed. Inferred ventilation index (IVI) was then calculated as the product of the average area of the peaks (mV x ms) and the number of peaks within 20 sec. At the end of the experiment (P110), animals were sacrificed and diaphragms collected for immunostaining.

### Immunofluorescence

#### Murine tissues

Following motor or respiratory evaluation, after cervical dislocation, murine SOD1^G93A^ SOL, EDL, diaphragms and Levator auris longus (LAL) muscles were dissected and fixed in 4% PFA (wt/vol) in PBS for 20 min at RT. SOL, EDL and diaphragms were further separated in bundles of 10-20 myofibers. Immunostaining of LAL muscles was performed using the whole-mount protocol (Negro *et al*., 2017b). After 20 min quenching in 50 mM NH_4_Cl in PBS, samples were saturated in blocking solution for 2h (15% goat serum, 2% BSA, 0.25% gelatine, 0.20% glycine, 0.5% Triton X-100 in PBS), and then incubated with specific primary antibodies in blocking solution for 72 h at 4°C. Axons and nerve terminals were labeled for VAMP-1 and NF. Tissues were then washed three times with PBS. Bundles were incubated at RT for 90 min in PBS with specific Alexa-fluor conjugates to identify postsynaptic nAChR. Images were collected with a Zeiss LSM 900 Confocal microscope equipped with a 40× HCX PL APO NA 1.4 oil immersion objective. To reduce cross talk, the laser excitation line, power intensity, and emission range were chosen based on the specific fluorophore of each sample.

#### Swine tissues

Diaphragms from hSOD1^G93A^ swines were cryo-sliced in 20 μm thick sections using Leica CM1520 cryostat after snap freezing. Slices were processed for immunostaining as described above.

#### Human tissues

Snap-frozen human muscle biopsies were fixed and split into bundles. Before incubation with the specific primary antibodies, bundles were incubated overnight at 4°C with α-BTx-555 in PBS with 0.5% BSA. Quenching and saturation were performed as previously described. Incubation with anti-CXCR4 antibody was prolonged for 4 d, followed by washes with PBS and a 3-h incubation at room temperature in dark conditions with secondary antibodies and Alexa-555- conjugated α-BTx diluted in blocking solution. Samples were then mounted under a stereomicroscope and imaged (Motanova *et al*, 2025).

### NMJ occupancy

Coloc Threshold plugin (Fiji) was used to measure overlap between presynaptic (VAMP-1) and postsynaptic (nAChRs) markers (namely NMJ occupancy). Percentages of denervated, partially innervated and innervated NMJs were obtained through Coloc threshold analysis measuring the Manders’ overlap coefficient (Dunn *et al*, 2011). Endplates were categorized based on occupancy percentage as follows: denervated (<40% overlap), innervated (>80% overlap), and partially denervated (40-80% overlap). This classification was adapted from similar methodologies previously applied to murine NMJs (Ang *et al*, 2022; Courtney *et al*, 2019). At least 6 fields of view and 50 NMJs were randomly selected and analysed for each muscle.

### MN survival

For the stereological analysis of spinal MN, the spinal cord was removed at cervical, thoracic and lumbar levels and post fixed in 4% PFA for 2 h. After cryoprotection overnight into a 30% sucrose solution in 0.1 M PBS buffer, the tissue was then embedded and frozen in cryostat medium (Killik, Bio-Optica, Milan, Italy). 50 μm thick, free-floating coronal sections of each spinal cord level were cut using a cryostat (Leica), then stored in an anti-freeze solution (30% ethylene glycol, 30% glycerol, 10% PB; 189 mM NaH_2_PO_4_; 192.5 mM NaOH; pH 7.4) at −20°C until being used. For Nissl staining, serial sections (1 every 400 μm, containing a representative series of the specific spinal tract) were washed in 0.1 M PBS (pH 7.4), then mounted on 4% gelatin-coated slides (Bio-Optica) and air-dried overnight. The following day, slices were hydrated in distilled water, immersed in 0.1% Cresyl violet acetate (Merck, C5042) for 8 min, dehydrated in ascending series of ethanol (95-100%) and cleared in xylene. Finally, the slides were coverslipped with Eukitt (Bio-Optica). MN stereological counts were performed along the tract C3-C5 for the cervical, T2-T5 for the thoracic, and L4-L5 for the lumbar segments. Only the nucleoli of multipolar neurons with an area ≥ 200 μm^2^ (attributable to αMNs) located in the spinal ventral horns at the proper position (Rexed lamina IX) for cervical, thoracic and lumbar traits were identified and counted (40x). The Optical Fractionator stereological technique was employed using a computer-assisted Nikon Eclipse E600 microscope equipped with the StereoInvestigator software (MicroBrightField Inc., Williston, VT, USA) (West *et al*, 1991).

Counts were performed using a 3 μm guard zone, 100 × 100 μm counting frame, and 150 × 150 μm scan grid. Ventral horn volume and cell numbers were obtained with NeuroExplorer (MicroBrightField). Cells were counted onscreen using a Nikon FDX-35 digital camera. Cell density was expressed as MNs/mm³ (Boido *et al*, 2014; Galbiati *et al*, 2023). Representative Nissl-stained images of cervical, thoracic, and lumbar ventral MNs were acquired with 10× and 20× objectives using the same camera.

### In vitro axonal elongation assay

Microfluidic chambers (MFC) were fabricated following established protocols (Park *et al*, 2006). Sterilized polydimethylsiloxane (PDMS) inserts (Dow Corning) were securely attached to 50-mm glass-bottomed WillCo dishes (IntraCel) using plasma cleaning. To prevent non-specific binding, the chambers were blocked overnight at 37°C with 0.8% BSA in PBS. For in vitro experiments, culture chambers were coated with Matrigel for 2 h at 37°C (Fornetti *et al*, 2022). On day 5, MN progenitors (from human iPSCs, see above) were dissociated and seeded into the proximal chamber at 3 × 10⁴ cells per device. After 30 min at 37°C to allow cell attachment, culture medium was added to all four reservoirs. On day 7, NUCC-390 was added to the distal chamber at 500 ng/mL and maintained for 5 d, and the medium changed every other day. On the fifth day, cells were fixed with 4% PFA in PBS at 4°C for 20 min, followed by immunofluorescence using standard protocol (Negro *et al*, 2018), and axonal elongation determined using NeuronJ plugin of Fiji software (Negro *et al*., 2017b).

### Assessment of NfL plasma levels

Mouse plasma samples were collected in 2 ml blood collection tubes and centrifuged at 5,000 x g for 5 min to obtain plasma. Neurofilament light chain (NfL) plasma levels were measured at baseline (P60) and at the end of NUCC-390 treatment period (P110) using a Simoa Nf-Light Advantage V2 kit (#104073) on the Quanterix SR-X^TM^ platform, according to the manufacturer’s protocol.

### Statistical analysis

Sample size was determined based on data collected by our laboratory or on published studies. The number of animals used/group and the number of muscle fibres/mouse for electrophysiological analysis and for imaging are reported in figure legends. Experiments were conducted under blinded conditions. No samples or animals were excluded from the analysis. To evaluate the impact of the CXCR4 agonist through functional and behavioural assays, animals were evenly distributed by sex, with male and female cohorts analyzed separately. GraphPad Prism software was used for all statistical analyses. Data normality was assessed using the Shapiro-Wilk test. Statistical significance was evaluated using analysis of variance (ANOVA) with Tukey post-test, Kruskal-Wallis non-parametric test, two-way ANOVA with Sidak post-test, mixed effect analysis test, and two-way ANOVA multiple comparisons. Data were considered statistically different when *p < 0.05, **p < 0.01, ***p < 0.001, ****p < 0.0001.

## Acknowledgements

This study was supported by:

Arisla Foundation: ‘Boosting REgeneration in ALS motor neurons by tArgeTing the peripHery’ (to MR)

Cariparo Foundation: ‘CXCR4: a marker of neurotransmission failure and a target for neuromuscular function recovery’ (to MR)

Stars Unipd: ‘CXCR4: A unifying target to FighT ALS from the pERiphery’ (to MP)

Italian Ministry of Health, Starting Grant Ricerca Finalizzata 2019: ‘Boosting peripheral nerve regeneration in ALS by CXCL12-CXCR4 axis’ (to SN) University of Padua (to MR and MP)

Italian Ministry of Defense, Project RIPANE (MONT_COMM18_03) (to CM)

We would like to thank Dr. Caterina Bendotti and Dr. James Sleigh for their critical reading of the manuscript and the fruitful discussion.

## Authors’ contributions

SN, CB, GDE, FF, GZ, MT performed the bench work. SN, CB, GDE performed motor function tests and relative analysis. FF, CB and SN performed the ventilation assay and relative analysis. MLM and AB managed mice colonies. RS and MB performed MN survival assessment. AR generated iPSC- derived hMNsFUS^P525L^ and IG and EF performed the related in vitro assays. VB and MB provided murine muscle samples, MF and LP analyzed blood NfL levels. The SOD1^G93A^ transgenic swine line was generated by RD and AP and clinically characterized by GC, whose muscle samples were provided by CT and CC. GS provided human muscle biopsies. AMe supervised electrophysiological assays. AF and AMa synthesized NUCC-390. MR, MP, CM and SN conceived the project and designed the experimental plan. MR, MP, CM and SN provided funding. MR wrote the manuscript with editing and intellectual input from MP and SN. All authors read and approved the final manuscript.

The authors declare that they have no conflict of interest

## Expanded Figure Legends

**Figure EV1.**
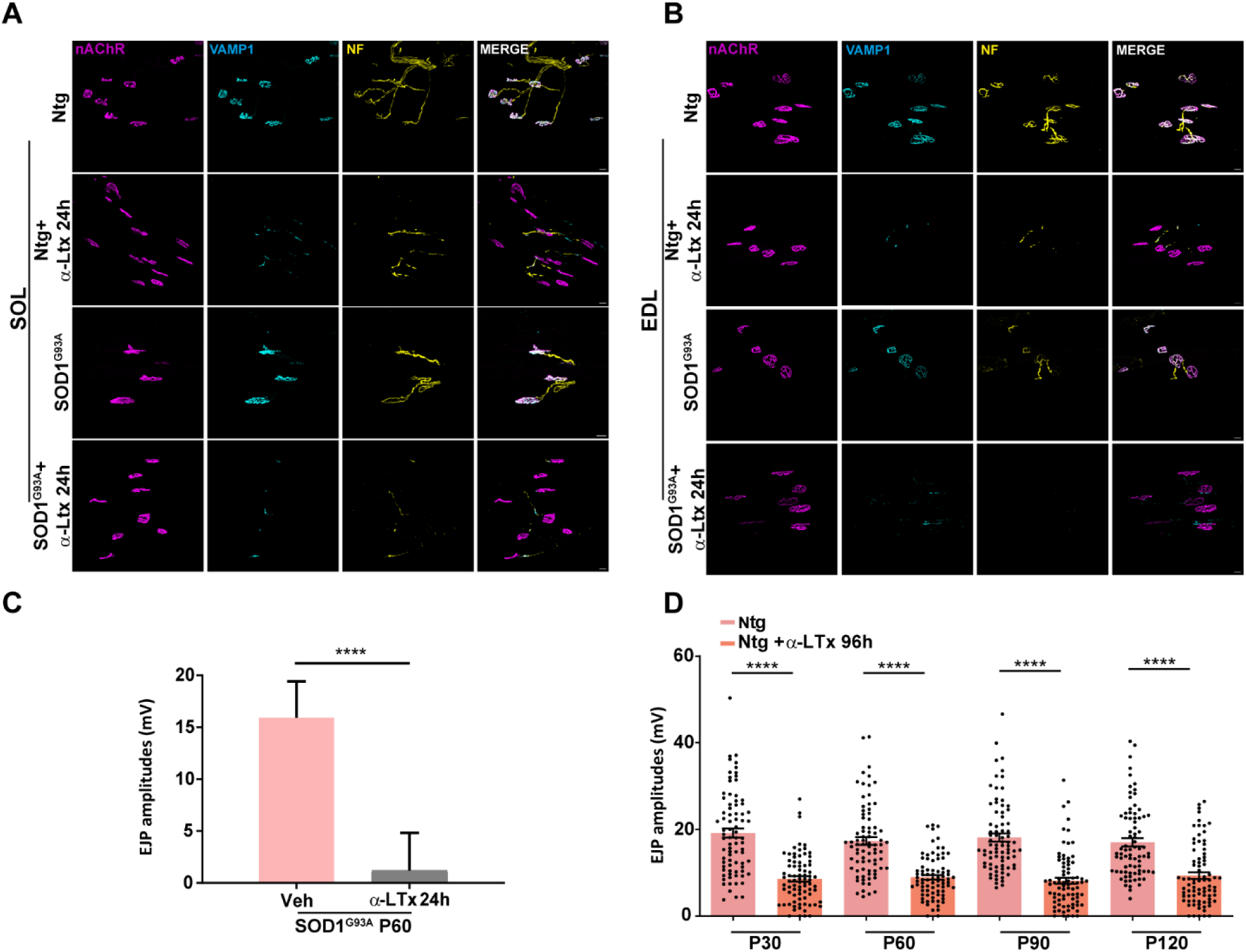
α-LTx is fully active in Ntg and SOD1^G93A^ mice. A-B Immunostaining of Ntg and SOD1^G93A^ NMJs (P60) 24 h from α-LTx injection in the SOL (A) and EDL (B). Presynaptic terminals are stained for VAMP1 (cyan) and NF (yellow), postsynaptic acetylcholine receptors are marked with α-BTx (nAChR, magenta). Scale bar: 20 µm. C EJP amplitudes recorded in 2 months-old SOD1^G93A^ 24 h from α-LTx injection. Box plots show median values. N=4, 20 fibers/mouse. Unpaired t-test **** P< 0,0001. D EJP amplitudes recorded in Ntg SOL before and 96 h from α-LTx injection performed at different disease stages (P30 to P120). Plots show median values and dots (black) single fiber distribution. N=4, 20 fibers/mouse. Kruskal Wallis test **** P<0,0001.

**Figure S2.**
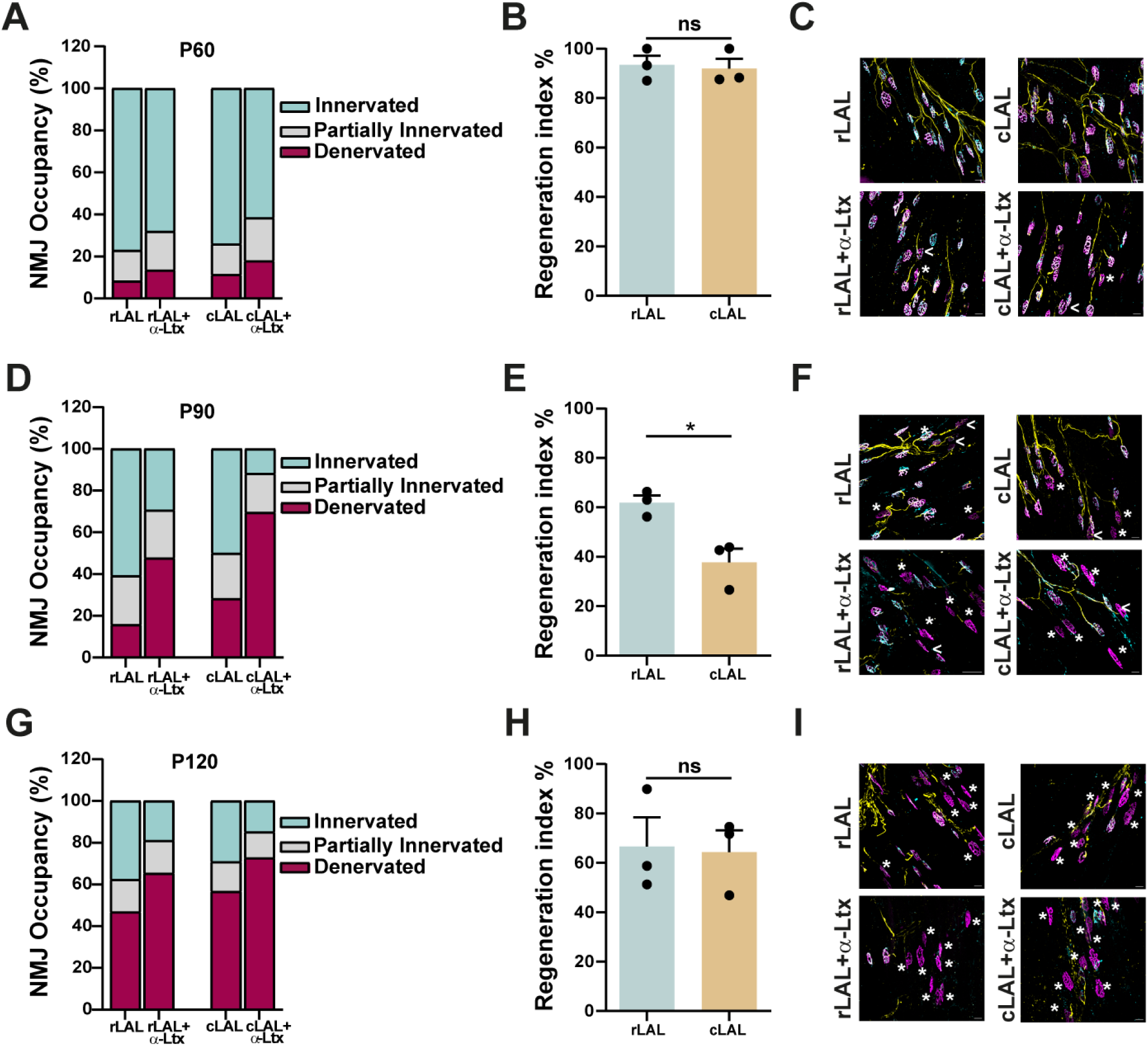
Assessment of NMJ regeneration in SOD1^G93A^ LAL during disease progression. A, D, G NMJ occupancy in rLAL and cLAL muscles at different disease stages (P60, P90, and P120, respectively), before and 96 h after α-LTx injection. NMJs are categorized as fully innervated, partially innervated or fully denervated based on the percentage of overlap between the presynaptic and postsynaptic compartments. B, E, H Regeneration index of SOD1^G93A^ rLAL and cLAL for each disease stage. Unpaired t test. *P=0,0188, ns=not significant. N=3, at least 80 NMJs analysed. C, F, I Representative immunostainings of rLAL and cLAL NMJs 96 h post α-LTx injection performed at different disease stages. Presynaptic nerve terminals are labelled for NF (yellow) and VAMP1 (cyan), postsynaptic acetylcholine receptors are marked with fluorescent α-BTx (nAChR, magenta). White arrowheads indicate partially innervated NMJs, while asterisks highlight denervated NMJs. Scale bar: 50 µm.

**Figure S3.**
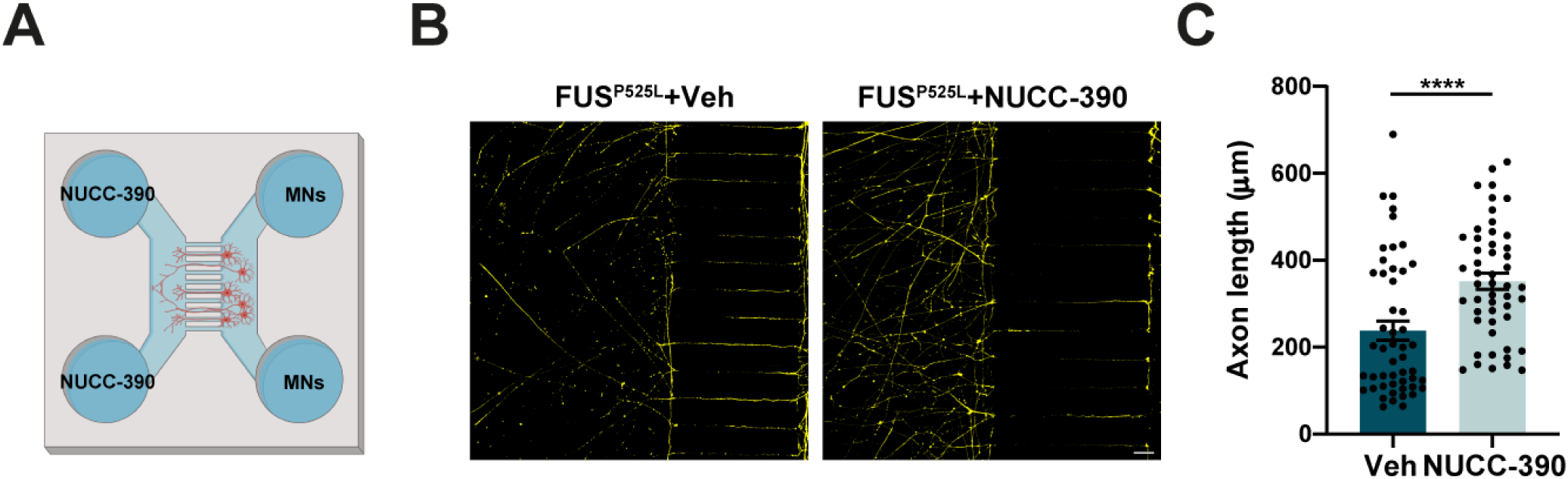
NUCC-390 promotes axonal elongation in hiPSC-derived ALS MNs. A Experimental design. Human iPSC-derived MNs carrying the pathogenic FUS^P525L^ mutation are grown on MFC. B Following NUCC-390 treatment for 5 d, cells are fixed and labelled for β3-tubulin (yellow). C Quantification of motor axon growth. Plots show median values and dots single-axon distribution. Axon length is expressed as µm. N=3, at least 15 axons/MCF, Mann Whitney test **** P<0,0001.

